# DRD2 *Taq*I A polymorphism in Eastern Uttar Pradesh population

**DOI:** 10.1101/783514

**Authors:** Amrita Chaudhary, Upendra Yadav, Pradeep Kumar, Vandana Rai

**Affiliations:** Human Molecular Genetics Laboratory, Department of Biotechnology, VBS Purvanchal University, Jaunpur-222003

**Keywords:** Dopamine D2 receptor, gene polymorphism, dopamine, PCR-RFLP, psychiatric disorders

## Abstract

Dopamine receptor D2 (DRD2) encoded by DRD2 gene, is located on chromosome 11q22-23. Dopamine plays the central role in motivation, cognition, and reward seeking behaviour. Its dysfunction is implicated in numerous neurological and psychiatric disorders including drug abuse, schizophrenia, ADHD etc. The *Taq*I A polymorphism is localized 9.8 kb downstream from DRD2 gene in exon 8 of protein kinase gene (ANKK1). It is a SNP demonstrated to cause Glutamate to Lysine substitution at 713 amino acid residue in putative binding domain of ANKK1. Due to the central role of dopamine in reward seeking behavior, DRD2 *Taq*I A loci is a suitable candidate for investigation of molecular basis of addiction. The aim of the present study is to evaluate the frequency of DRD2 *Taq*I A polymorphism in Eastern Uttar Pradesh population. 3ml blood samples were collected from 50 individuals randomly selected from Eastern UP. Written informed consent along with profile detail was taken from each subject prior to blood sample collection. DRD2 *Taq*I A polymorphism analysis was done by PCR-RFLP method. Genomic DNA was extracted from each collected blood samples and amplified using DRD2 *Taq*1 region specific primers. PCR amplification produced 310bp long amplicon which was digested with *Taq* I enzyme for polymorphism analysis. In case of A2 allele, *Taq*1 enzyme cleaved 310bp long fragment into two fragments of 180bp and 130bp. In case of A1 allele, a C to T substitution demolished the restriction site of *Taq*1, so amplicon of A1 allele remained uncut. In total 50 sample analyzed in present study, A2/A2, A2/A1 and A1/A1 genotype were found in 12, 32 and 06 samples respectively. The genotypic frequencies of mutant homozygous (A1/A1) is 0.12, heterozygous (A2/A1) is 0.64 and normal homozygous (A2/A2) is 0.24. The allelic frequency of A1 is 0.44 and of A2 is 0.56. In conclusion, the results of present study suggests that in *Taq*I A polymorphism of DRD2 gene, the frequency of allele A2 is higher than that of A1 allele in population of Eastern Uttar Pradesh.

## Introduction

One of the most important system intervening reward mechanisms is considered to be dopaminergic pathways. Dopaminergic neurons are present in VTA of midbrain, projected into nucleus accumbens and ventral striatum [1] All the genes, involved in regulating the assembly of this system in brain is of great interest and can be a suitable candidate for investigation of molecular basis of addiction and several other psychiatric disorders [2]. Among these, the gene of interest that effect the dopaminergic neurotransmission is the Dopamine D2 receptor (DRD2) gene that is located at chromosome 11q22-23 encoding a G-Protein coupled receptor (Gi - inhibitory G protein) in post synaptic neurons [3] performing dual function of inhibitory auto receptor and a post synaptic receptors [4].

Several polymorphisms are reported in DRD2 gene (*Taq*1 B, *Taq*1D, −141 Ins/Del, Ser-Cys(S311C) but *Taq*1 A polymorphism is well studied in different psychiatric disorders including schizophrenia [5], depression [6], bipolar disorder [7], ADHD [8], and PTSD [9] and drug abuse [3]. *Taq*I A is SNP (rs1800497) located 9.8 kb downstream of DRD2 gene within exon 8 of functionally unrelated neighbouring gene, Ankyrin repeat and kinase domain containing-1(ANNK1). It causes Glutamate to Lysine substitution at 713 amino acid residue in putative binding domain of ANNK1 [10] with two alleles attributed as A1 and A2. This polymorphism leads to alteration of the activity of promoter region of DRD2 gene and also the expression of D2-type receptors [11]. Individuals with A1A1 genotype have approximately 49% reduced number of dopamine receptors than to those without A1. Less number of DRD2 receptors in nucleus accumbens and striatum increases craving for alcohol. Hence, the aim of present study is to determine the frequency of DRD2 *Taq*IA polymorphism in Eastern Uttar Pradesh population.

## Materials and Methods

All 50 participants were recruited from Eastern Uttar Pradesh (Jaunpur) population, informed written consent was taken from each participants. Prior blood sample collection clearance certificate was taken from the Institutional Ethics Committee of VBS Purvanchal University, Jaunpur.

### Genotyping

3 ml blood was drawn from each individual and genomic DNA was isolated by the method of Bartlett and White [12]. Polymerase chain reaction (PCR) based genotyping was done by using gene specific primers according to the method of Grandy et al. [13]. The reaction conditions were enlisted in table 1. Restriction *Taq*1 enzyme digested products were separated on 2% agarose gel with 100 bp marker.

**Table 1.**
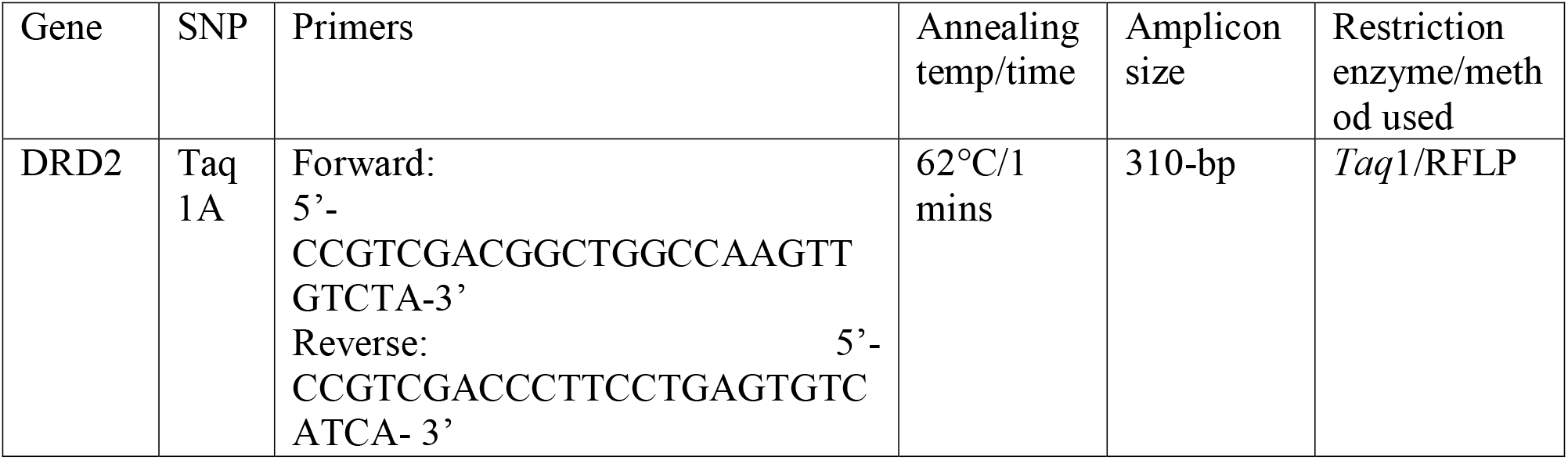
Representing gene, PCR primers, annealing temperature, time, and restriction enzyme used.

### Statistical Analysis

Allele frequency was determined by gene counting method.

## Results and Discussion

PCR amplification produced 310 bp long amplicon, which after digestion with *Taq*I produced 180bp and 130 bp long fragments in case of normal C allele (A2) and mutant T allele (A1) remain uncut after *Taq*I digestion (Fig. 1 and 2).

**Fig. 1.**
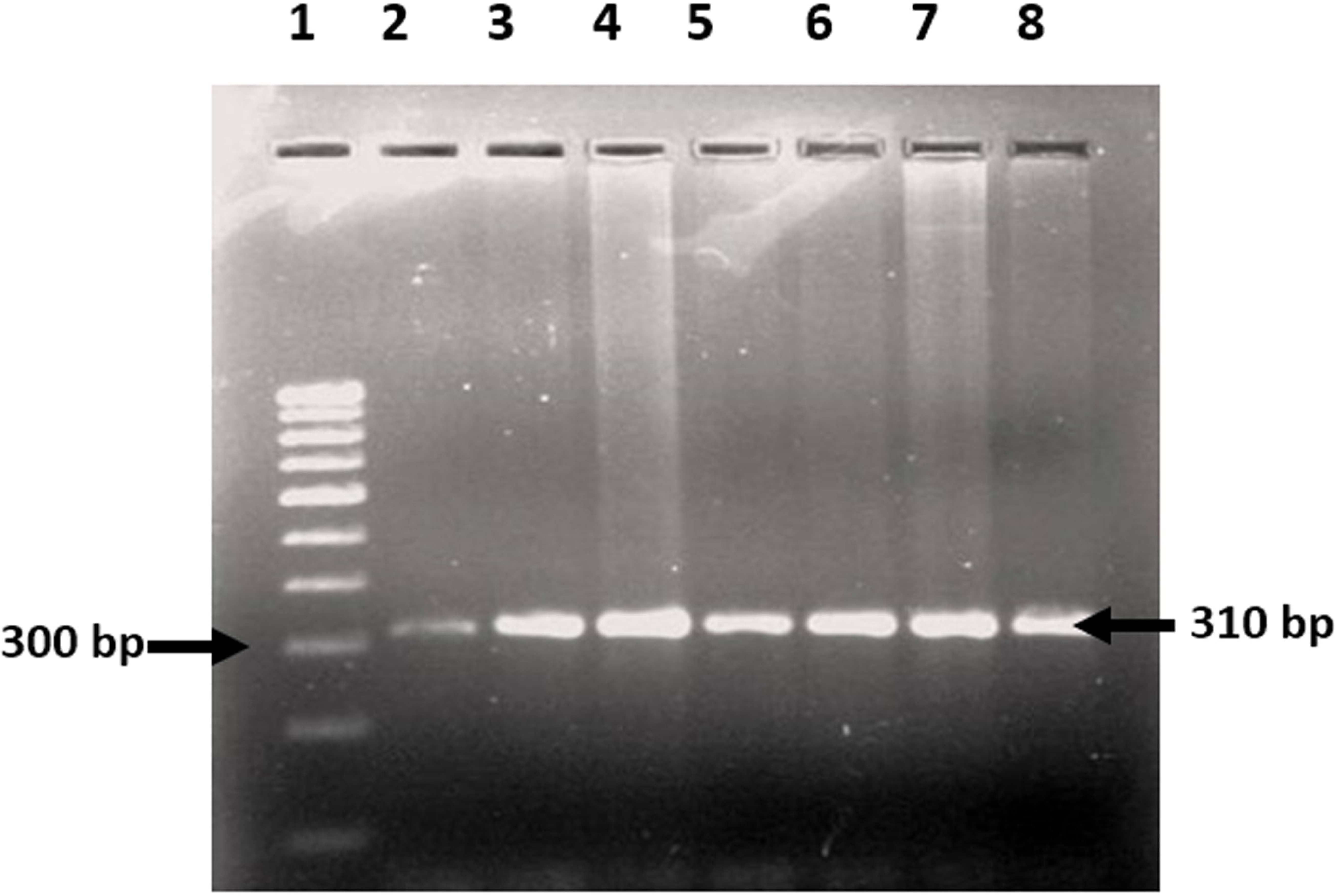
Agarose gel showing amplicon of 310 bp and 100-bp ladder in lane 1

**Fig. 2.**
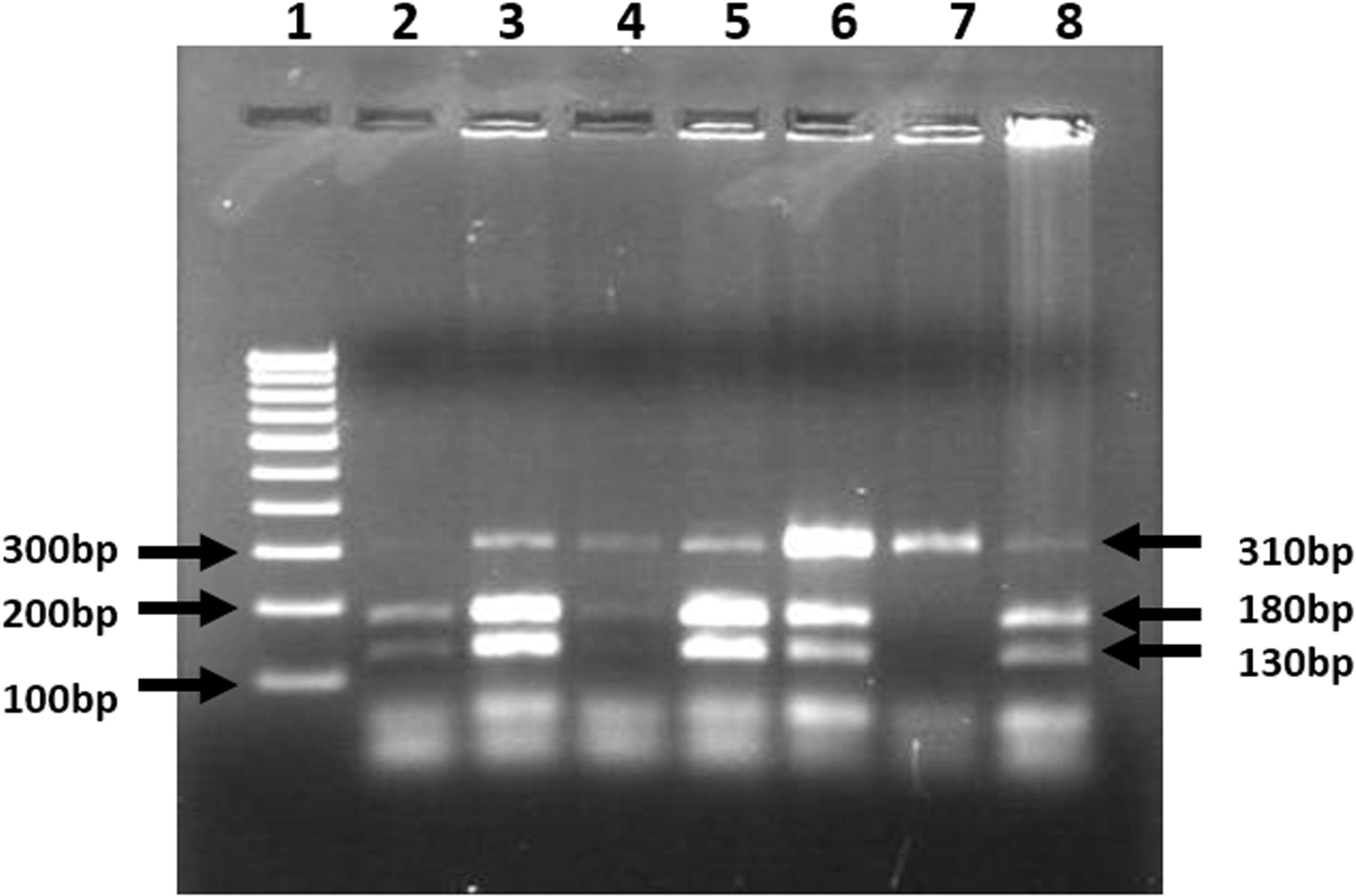
Restriction digestion of amplicon by *Taq*I with 100 bp marker in lane 1.

In total 50 samples analyzed, normal homozygous genotype was found in 12, heterozygous genotype was observed in 32 and mutant homozygous genotype was observed in 6 individuals. The C and T allele were found as 0.56 and 0.44 respectively (Table 2).

**Table 2.**
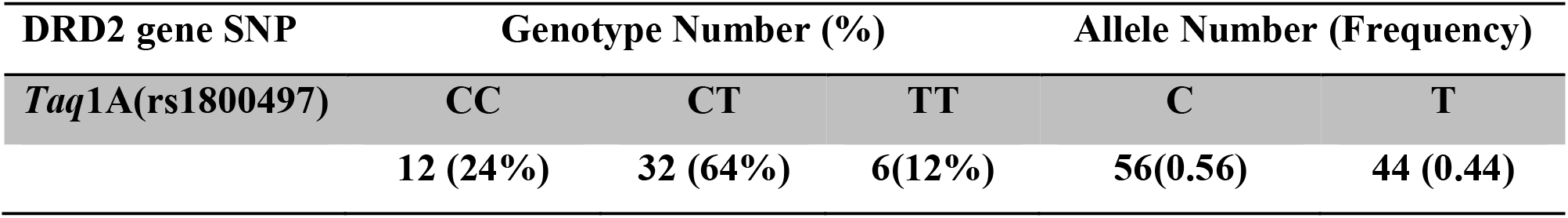
Genotype and allele frequency distribution in studied subjects.

Four alleles (genetic variants) are reported at *Taq1*A locus referred to as A1 A2, A3 and A4. Where A2 is the most common form and A3 and A4 are the rare variants [14]. DRD2gene *Taq*1A polymorphism was very well studied in world’s different populations [3, 5, 9, 15, 16] as well as in different Indian regions [17–27]. Various studies reported presence of A1 allele is significantly higher in certain drug of abuse such as alcoholism and other behavioural characteristics as well [28]. Here our report suggests that there is higher frequency of A2 allele in Jaunpur population. Our study is co-related with numerous studies from India where they also reported higher frequency of A2 allele in control population. Juyal et al. [26] studied South and North Indian population, worked on Parkinsons disease reported frequency of A2 allele to be 0.66 and A1 allele to be 0.33 in their control group. One other study of Kumudini et al. [29] studied South Indian population also found nearly the same result. They reported 0.68, A2 allele frequency and 0.31 A1 allele frequencies. However, few reports are also available that shows the higher frequency of A1 allele in control group. The report of Vijayan et al. [25] indicates the A2 allelic frequency is 0.34 and A1 allele is 0.65. Various international studies also report the higher prevalence of A2 allele in their control group. Alfimova et al. [5] stated A2 allele frequency to be of 0.80 and A1 allele to be of 0.19 in Moscow population. Another study on PTSD of Voisey et al. [15] of Australia studied control Caucasians reported A2 allele frequency 0.82 and A1 allelic frequency 0.17. Despite contradictory results were indicated by a study of China by Lee et al. [16] worked on Tourette syndrome explored control group and reported 0.47 A2 allele frequency and 0.52 A1 allele frequency. Hence, these findings showing the variations in the allelic frequency suggest that prevalence of higher A2 allelic frequency can be population specific.

## References

1. Matosic, A., Marusic, S., Vidrih, B., Kovak-Mufic, A., Cicin-Sain, L.: Neurobiological Bases of Alcohol Addiction. Acta Clin Croat 55(1), 134–50 (2016).

2. Ma, Y., Yuan, W., Jiang, X, Cui, W.Y., Li, M.D.: Updated findings of the association and functional studies of DRD2/ANKK1 variants with addictions. Mol Neurobiol 51(1), 281–99 (2015).

3. Panduro, A., Ramos-Lopez, O., Campollo, O., Zepeda-Carrillo, E.A., Gonzalez-Aldaco, K., Torres-Valadez, R., Roman, S.: High frequency of the DRD2/ANKK1 A1 allele in Mexican Native Amerindians and Mestizos and its association with alcohol consumption. Drug Alcohol Depend 172, 66–72 (2017).

4. Tunbridge, E. M., Narajos, M., Harrison, C. H., Beresford, C., Cipriani, A., & Harrison, P. J.: Which Dopamine Polymorphisms Are Functional? Systematic Review and Meta-analysis of COMT, DAT, DBH, DDC, DRD1-5, MAOA, MAOB, TH, VMAT1, and VMAT2. Biological Psychiatry. (2019). doi:10.1016/j.biopsych.2019.05.014

5. Alfimova, M. V., Golimbet, V. E., Korovaitseva, G. I., Lezheiko, T. V., Tikhonov, D. V., Ganisheva, T. K., Berezin, N.B., Snegireva, A.A., Shemyakina, T. K.: The Role of the Interaction between the NMDA and Dopamine Receptor Genes in Impaired Recognition of Emotional Expression in Schizophrenia. Neuroscience and Behavioral Physiology 49(1), 153–158 (2019).

6. Hayden, E. P., Klein, D. N., Dougherty, L. R., Olino, T. M., Laptook, R. S., Dyson, M. W., Bufferd, S.J., Durbin, E., Sheikh, H.I., Singh, S. M.: The dopamine D2 receptor gene and depressive and anxious symptoms in childhood: associations and evidence for gene–environment correlation and gene–environment interaction. Psychiatric Genetics 20(6), 304–310 (2010).

7. Hu, M.C., Lee, S.Y., Wang, T.Y., Chang, Y.H., Chen, S.L., Chen, S.H., Chu, C.H., Wang, C.L., Lee, I.H., Chen, P.S., Yang, Y.K., Lu, R.B.: Interaction of DRD2TaqI, COMT, and ALDH2 genes associated with bipolar II disorder comorbid with anxiety disorders in Han Chinese in Taiwan. Metab Brain Dis 30(3), 755–65 (2015).

8. Sery, O., Drtilkova, I., Theiner, P., Pitelova, R., Staif, R., Znojil, V., Lochman, J., Didden, W.: Polymorphism of DRD2 gene and ADHD. Neuro Endocrinol Lett 27(1-2), 236–40 (2006).

9. Xiao, Y., Liu, D., Liu, K., Wu, C., Zhang, H., Niu, Y., & Jiang, X.: Association of DRD2, 5-HTTLPR, and 5-HTTVNTR Gene Polymorphisms With Posttraumatic Stress Disorder in Tibetan Adolescents: A Case–Control Study. Biological Research For Nursing 21(3), 286–295 (2019).

10. Pan, Y.Q., Qiao, L., Xue, X.D., Fu, J.H.: Association between ANKK1 (rs1800497) polymorphism of DRD2 gene and attention deficit hyperactivity disorder: a meta-analysis. Neurosci Lett 590, 101–5 (2015).

11. Marinho, V., Oliveira, T., Bandeira, J., Pinto, G.R., Gomes, A., Lima, V., Magalhaes, F., Rocha, K., Ayres, C., Carvalho, V., Velasques, B., Ribeiro, P., Orsini, M., Bastos, V.H., Gupta, D., Teixeira, S.: Genetic influence alters the brain synchronism in perception and timing. J Biomed Sci 25(1), 61 (2018).

12. Bartlett, J.M., and White, A., Extraction of DNA from blood. In: Bartlett, J.M., Stirling, D., (eds.) Methods in Molecular Biology. PCR Protocols. 2^nd^ ed., vol. 226. Totowa, NJ: Humana Press Inc. (2003).

13. Grandy, D.K., Zhang, Y., Civelli, O.: PCR detection of the TaqA RFLP at the DRD2 locus. Hum Mol Genet 2(12), 2197 (1993).

14. Blum, K., Cull, J. G., Braverman, E. R., & Comings D. E.: Reward deficiency syndrome. American Scientist 84(2), 132–145 (1996a).

15. Voisey, J., Swagell, C.D., Hughes, I.P., Morris, C.P., van Daal, A., Noble, E.P., Kann, B., Heslop, K.A., Young, R.M., Lawford, B.R.: The DRD2 gene 957C>T polymorphism is associated with posttraumatic stress disorder in war veterans. Depress Anxiety 26(1), 28–33 (2009).

16. Lee, C.C., Chou, I.C., Tsai, C.H., Wang, T.R., Li, T.C., Tsai, F.J.: Dopamine receptor D2 gene polymorphisms are associated in Taiwanese children with Tourette syndrome. Pediatr Neurol 33(4), 272–6 (2005).

17. Kaur, G., Chavan, B.S., Gupta, D., Sinhmar, V., Prasad, R., Tripathi, A., Garg, P.D., Gupta, R., Khurana, H., Gautam, S., Margoob, M.A., Aneja, J.: An association study of dopaminergic (DRD2) and serotoninergic (5-HT2) gene polymorphism and schizophrenia in a North Indian population. Asian J Psychiatry 39, 178–184 (2019).

18. Roy, S., Pal, P., Ghosh, S., Bhattacharya, S., Das, S.K., Gangopadhyay, P.K., Bavdekar, A., Ray, K., Sengupta, M., Ray, J.: Potential Role of Brain-Derived Neurotrophic Factor and Dopamine Receptor D2 Gene Variants as Modifiers for the Susceptibility and Clinical Course of Wilson’s Disease. Neuromolecular Med 20(3), 401–408 (2018).

19. Quraishi, R., Jain, R., Mishra, A.K., Ambekar, A.: Association of ankyrin repeats & kinase domain containing 1 (ANKK1) gene polymorphism with co-morbid alcohol & nicotine dependence: A pilot study from a tertiary care treatment centre in north India. Indian J Med Res 145(1), 33–38 (2017).

20. Suraj Singh, H., Ghosh, P.K., Saraswathy, K.N.: DRD2 and ANKK1 gene polymorphisms and alcohol dependence: a case-control study among a Mendelian population of East Asian ancestry. Alcohol Alcohol 48(4), 409–14 (2013).

21. Bhaskar, L.V., Thangaraj, K., Non, A.L., Singh, L., Rao, V.R.: Population-based case-control study of DRD2 gene polymorphisms and alcoholism. J Addict Dis 29(4), 475–80 (2010).

22. Prasad, P., Ambekar, A., Vaswani, M.: Dopamine D2 receptor polymorphisms and susceptibility to alcohol dependence in Indian males: a preliminary study. BMC Med Genet 11, 24 (2010).

23. Srivastava, V., Deshpande, S.N., Thelma, B.K.: Dopaminergic pathway gene polymorphisms and genetic susceptibility to schizophrenia among north Indians. Neuropsychobiology 61(2), 64–70 (2010).

24. Prasad, P., Kumar, K.M., Ammini, A.C., Gupta, A., Gupta, R., Thelma, B.K.: Association of dopaminergic pathway gene polymorphisms with chronic renal insufficiency among Asian Indians with type-2 diabetes. BMC Genet 9, 26 (2008).

25. Vijayan, N.N., Bhaskaran, S., Koshy, L.V., Natarajan, C., Srinivas, L., Nair, C.M., Allencherry, P.M., Banerjee, M.: Association of dopamine receptor polymorphisms with schizophrenia and antipsychotic response in a South Indian population. Behav Brain Funct 3, 34 (2007).

26. Juyal, R.C., Das, M., Punia, S., Behari, M., Nainwal, G., Singh, S., Swaminath, P.V., Govindappa, S.T., Jayaram, S., Muthane, U.B., Thelma, B.K.: Genetic susceptibility to Parkinson’s disease among South and North Indians: I. Role of polymorphisms in dopamine receptor and transporter genes and association of DRD4 120-bp duplication marker. Neurogenetics 7(4), 223–9 (2006).

27. Shaikh, K.J., Naveen, D., Sherrin, T., Murthy, A., Thennarasu, K., Anand, A., Benegal, V., Jain, S.: Polymorphisms at the DRD2 locus in early-ons et al. ohol dependence in the Indian population. Addict Biol 6(4), 331–335 (2001).

28. Blum, K., Sheridan, P.J., Wood, R.C., Braverman, E.R., Chen, T.J., Cull, J.G., Comings, D.E.: The D2 dopamine receptor gene as a determinant of reward deficiency syndrome. J R Soc Med 89(7), 396–400 (1996b).

29. Kumudini, N., Umai, A., Devi, Y.P., Naushad, S.M., Mridula, R., Borgohain, R., Kutala, V.K.: Impact of COMT H108L, MAOB int 13 A>G and DRD2 haplotype on the susceptibility to Parkinson’s disease in South Indian subjects. Indian J Biochem Biophys 50(5), 436–41 (2013).

